# A Novel 3D Co-culture Platform for Integrating Tissue Interfaces for Tumor Growth, Migration and Therapeutic Sensitivity: “PP-3D-S”

**DOI:** 10.1101/2021.10.14.464428

**Authors:** Mansoureh Mohseni Garakani, Pouyan Ahangar, Sean Watson, Bernard Nisol, Michael R. Wertheimer, Derek H. Rosenzweig, Abdellah Ajji

## Abstract

Metastatic cancers can be highly heterogeneous, show large patient variability and are typically hard to treat due to chemoresistance. Personalized therapies are therefore needed to suppress tumor growth and enhance patient’s quality of life. Identifying appropriate patient-specific therapies remains a challenge though, due mainly to non-physiological *in vitro* culture systems. Therefore, more complex and physiological *in vitro* human cancer microenvironment tools could drastically aid in development of new therapies. We developed a plasma-modified, electro-spun 3D scaffold (PP-3D-S) that can mimic the human cancer microenvironment for customized-cancer therapeutic screening. The PP-3D-S were characterized for optimal plasma-modifying treatment and scaffolds morphology including fiber diameter and pore size. PP-3D-S was then seeded with human fibroblasts to mimic a stromal tissue layer; cell adhesion on plasma-modified poly (lactic acid), PLA, electrospun mats vastly exceeded that on untreated controls. The cell-seeded scaffolds were then overlaid with alginate/gelatin-based hydrogel embedded with MDA-MB231 human breast cancer cells, representing a tumor-tissue interface. Among three different plasma treatments, we found that NH^3^ plasma promoted the most tumor cell migration to the scaffold surfaces after 7 days of culture. For all treated and non-treated mats, we observed a significant difference in tumor cell migration between small-sized and either medium- or large-sized scaffolds. In addition, we found that the PP-3D-S was highly comparable to the standard Matrigel® migration assays in two different sets of doxorubicin screening experiments, where 75% reduction in migration was achieved with 0.5μM doxorubicin for both systems. Taken together, our data indicate that PP-3D-S is an effective, low-cost, and easy-to-use alternate 3D tumor migration model which may be suitable as a physiological drug screening tool for personalized medicine against metastatic cancers.

## 1. Introduction

According to a recent report from Canadian Cancer Statistics [1], close to 50% of Canadians will develop cancer in their lifetime, and nearly half of these will die of the disease. Based on this report, 220,400 Canadians were diagnosed with cancer and 82,100 were estimated to have died just in 2019. The most frequently diagnosed cancers were predicted to be lung, breast, colorectal and prostate cancers, accounting for 48% of all cases in 2019. Therefore, the importance of research activities in cancer disease and its cure is obviously crucial. In oncology, treatments such as surgery, radio-therapy and chemotherapy frequently lead to severe side effects, greatly decreasing the quality of life of patients [2]. Development of novel therapeutic strategies is therefore needed to treat cancers more effectively while also improving overall patient outcomes. However, developing new therapies is a lengthy and costly path, with 75-90% of new drug candidates failing to pass phase 3 clinical trials [3-5]. Biofabrication and 3D culture technology has been gaining a lot of traction and may be quite suitable toward developing new cancer treatments.

Biofabrication, a rapidly growing technology mainly stimulated by advances in 3D fabrication techniques, is defined as the production of a customized 3D construct or complex biological products in which living cells, proteins, bioactive molecules, extracellular matrices, and biomaterials are considered as the building blocks to manufacture the model. Constructs developed by this strategy have the potential to recapitulate the complexity and heterogeneity of tissues and organs, so will be able to be applied for tissue regeneration and regenerative medicine and / or used as *in vitro* 3D models for drug discovery [6-8]. In cancer studies, biofabrication can be used to recreate or model various tissues using human cells, thereby presenting a physiological microenvironment such as the tumor-tissue interface. By developing artificial cancer tissue models, it will be possible to postpone the above-named drastic therapies, and to enable oncologists to predict and monitor how the disease progresses. It is noteworthy that microenvironment of neoplasia, specifically stromal-epithelial interactions which are critical and essential for cancer therapy, are still underexplored due to insufficient understanding of stromal-epithelial crosstalk [9, 10]. Furthermore, cell culture on two-dimensional (2D) flat polystyrene culture dishes does not realistically portray the behaviour of living cells in their normal three-dimensional (3D) environment of natural tissues [11]. Different types of cancers may have variations in complexity of tumor microenvironments, and it is challenging to mimic intercellular interaction *in vitro* [12, 13]; to do so requires realistic and physiological tissue models. The most common issue with 2D cell culture is that it does not reproduce complexities of cell-extracellular matrix (ECM) interactions *in vivo*. Standard 2D culture often fails to recapitulate interactions between cancer epithelial cells and stromal compartment, which play a crucial role in tumorigenesis and progression. This has directly led to a rise in developing 3D culture and co-culture models of cancer microenvironments [13, 14]. Cancer cells in 3D systems better mimic *in vivo* tumor growth compared with 2D culture [11-14]. This allows for long-term investigation and screening of novel therapeutics for anti-cancer efficacy using either cell lines or patient derived materials. Thus, the 3D cancer tissue model is more promising for high-throughput screening of anticancer drugs [9, 12, 15]. Numerous 3D culture products have been developed in recent years to model the tumor microenvironment, namely scaffold-free and scaffold-based culture systems including hydrogels, sol gels, ceramic or polymeric 3D-printed scaffolds, expanded polystyrene supports, permeable membranes, and electro-spun nanofiber layers, to name but a few [16].

Natural components of the ECM such as collagen, fibronectin and hyaluronic acid, as well as alternates such as gelatin, and alginate, have been developed as hydrogels to provide structural support for cell interactions in 3D culture systems. They have been the most commonly used materials due to their inherent cytocompatibility, intrinsic cell adhesion properties and capability of being remodelled by cells [14, 17, 18]. Commercially available basement membrane extract (BME), a gelatinous protein mixture secreted by certain murine sarcoma cells, resembles the complex ECM environment found in many tissues [19, 20]. Indeed, the heterogeneous composition of those ECM structural proteins includes laminin, collagen and proteoglycans, among others, which present cultured cells with the adhesive peptide sequences encountered in their natural environment. Several suppliers offer this product, for example under the trade names Matrigel^®^ or Cultrex^®^, for purposes of research in cancer biology (e.g. cell-migratory behavior as a model of tumor cell), and for use by pharmaceutical scientists to screen potential anti-cancer molecules [21]. However, these products are expensive, require multiple time-consuming steps for implementation, exhibit batch-to-batch variability, have limited mechanical strength and uncontrolled degradation. Furthermore, they do not truly mimic the biophysical and biochemical networks found in native ECM *in vivo* [20, 22]. Data gathered from tumor cell growth using such materials may not present optimal results compared to *in vivo* animal studies or patient data.

To overcome the above-listed limitations, other approaches of 3D tumor models such as spheroids [23-26] or organoids [27] have also been trending in cancer research for over 2 decades. There are four different types of 3D spherical, *in vitro* tumor models [28] Including multicellular tumor spheroids developed by the hanging drop technique (HDT), tumorospheres produced from cancer stem cells, tissue-derived tumorospheres created from partially digested cancer tissues, and organotypic multicellular spheroids made from cut sections of tumors. These models have proven valuable in identifying differences in drug responses to new and existing therapeutics, opening doors for personalized medicine [29]. However, most organoid and spheroid models lack the cellular, biomolecular, and biomechanical heterogeneity of the tumor microenvironment *in vivo*. Therefore, new pre-clinical screening platforms that better mimic the complex tumor physiological microenvironment are necessary to enhance regulatory approval and clinical translation.

Metastatic or secondary cancer is a type that has spread from the original so-called primary tumor to another part of the body [30, 31]. Bone tissue is a frequent site to be affected by metastatic cancer, termed as bone metastasis, and tumor arising from the breast, prostate, lung, and kidney are the most common sources [32]. Several *in vitro* 3D models of bone metastasis have emerged over the past 10-15 years such as 3D explant co-culture [33], 3D printed composite scaffolds [34, 35] and electrospun mesh co-culture [36]. These models are showing promise in anti-cancer drug development and for better understanding of pathophysiology. Solid scaffolds (porous and/or fibrous) made of synthetic materials such as polyethylene glycol (PEG), poly(lactide-co glycolide) (PLG), polycaprolactone (PCL) and poly(lactic acid) (PLA) have advantages of reproducibility, being inert, with “tunable” biodegradability, and more specifically, control on tuning mechanical properties and chemical composition for realistically mimicking the ECM of tumors. The most powerful method to produce fibrous scaffolds based on synthetic materials is *electrospinning*; there already exists a substantial literature regarding their biomedical applications [37-42]. Indeed, electrospun mats are particularly attractive because they can be made from various soft or stiff materials and can have tunable mechanical properties, porosity and fiber size, which allow for mimicking of natural tissues with various extracellular matrices [43, 44].

As stated earlier in this text, microenvironment of neoplasia, specifically stromal-epithelial cells interactions which have a critical and essential role in tumorigenesis and progression, are still underexplored due to insufficient understanding of stromal-epithelial crosstalk. Most 3D models are generated only from epithelial tumor cells; obviously, this cannot elucidate the molecular underpinnings of stromal-epithelial interactions. Therefore, 3D structures that contain both stromal and epithelial tumor cells constitute a clear and major improvement. Considering earlier-mentioned gaps in the literature, we aim to develop and perfect a 3D interface model comprised of both stromal and epithelial cancer cells, one that truly mimics the microenvironment of human cancers. Referred to hereafter as ***PP-3D-S*** (for **P**lasma-**P**olymer coated, electro-spun **3D S**caffold), this novel 3D co-culture model based on electrospinning is customizable, reproducible and closely resembles *in vivo* tissue matrix structures. It is also more cost-effective, versatile and easier to use than the above-mentioned ECM-derived commercial materials. As reported below, our group employs *PP-3D-S* to generate a 3D migration and invasion model resembling tumor microenvironment, to study tumor interface and metastasis. The invention can also deal with cancer therapies, by enabling one to screen anti-cancer therapeutics in a high-throughput manner. It is based on fabrication of 3D electrospun fibrous scaffolds with different materials and morphologies, followed by plasma functionalization of the fibre surfaces with appropriate functional groups or by coating them with thin bio-active films deposited by plasma polymerization. Stromal cells generally found in most tissues, like fibroblasts, can thereafter readily be seeded onto plasma-treated electrospun poly(lactic acid), PLA, for example; in this work, the mat is then overlaid with an alginate/gelatin-based hydrogel layer, pre-seeded with MDA-MB-231 aggressive breast cancer cell line. We hypothesize that plasma-bioactivated 3D scaffolds greatly improve cell adhesion and growth and stimulate invasive tumor cell migration through judicious choices of electrospun polymer scaffold and appropriate plasma treatments. We start by characterizing the materials, surface properties and morphological aspects, then followed by biological investigations such as cancer cell invasion and metastasis after interfacing scaffold and hydrogel.

## 2. Experimental Section

### 2.1. Preparation of plasma-treated electrospun nanofibrous scaffold

Figure 1 schematically portrays the various fabrication steps that characterize the preparation of the PP-3D-S plasma-treated electrospun nanofibrous scaffold technology being described in this present article.

**Figure 1.**
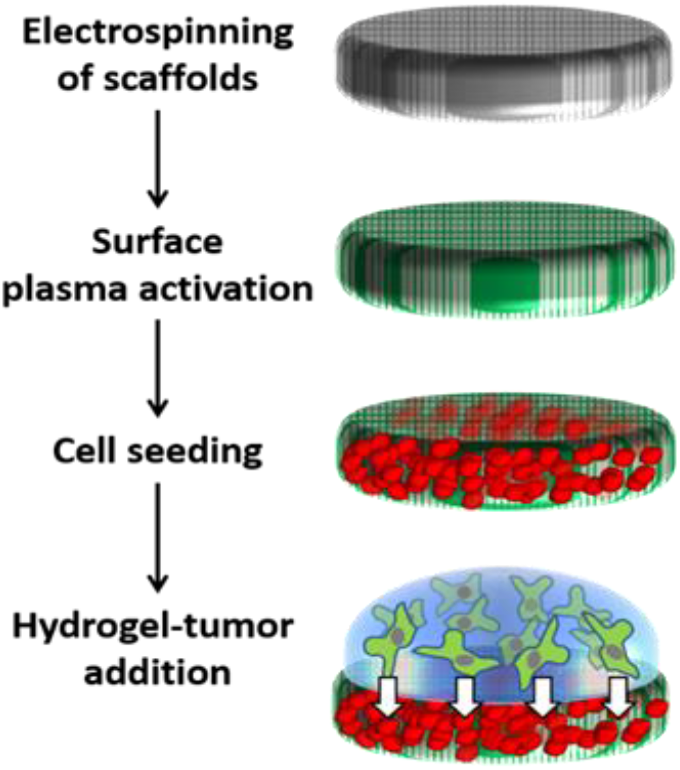
Schematic diagram of the main steps that comprise the current innovative methodology. Each of these is elaborated in successive sub-sections of the text in much greater detail.

#### 2.2.1 Fabrication of nanofibrous mat: electrospinning

As the first step, electrospinning was used to fabricate poly (lactic acid), PLA (NatureWorks 4032D, density of 1.24 g/cc) randomly-oriented nanofibrous scaffolds with finely-tuned morphology, in three different sizes of fiber diameters, hereafter designated “Small”, “Medium” and “Large”. Briefly, to produce a “Medium” PLA mat, 16 wt.% polymer solution was prepared by dissolving PLA pellets in 2,2,2 trifluoroethanol, TFE, (M=100.04 g/mol, Merck) solvent and stirring for 24h. Using a syringe pump placed in a chamber with a controlled temperature (21-24 ^0^C) and relative humidity of about 45%-50%, 10 ml of polymer solution (depending on the desired final mat thickness, which was intended to be about 250 μm) was electrospun with a flow rate of 1.6 ml/hr. The distance of the grounded needle tip (21G) and a rotating collector (25 rpm) was set at 15 cm, and the applied DC voltage of 20 kV between the needle tip and the rotating mandrel was provided by a stable DC high-voltage (HV) power supply. Nanofiber filaments were collected on the rotating metal drum, onto which an aluminum foil had been wrapped. The electrospun mats on the foil were then placed in ambient air to evaporate residual solvent for ca. 3 days, then gently detached, cut into smaller pieces, and stored in a desiccator for subsequent use. Two additional types of different medium-sized electro-spun polymer scaffolds, namely polyurethane, PU (MDI-polyester/polyether polyurethane, CAS 68084-39-9) and polycaprolactone, PCL (Sigma-Aldrich, Mn = 80 000) were also fabricated, to examine how versatile the methodology might be. For the case of PU a 1:1 mixture of tetrahydrofuran, THF, and N,N dimethylformamide, DMF, (both from Sigma) was used as solvent, while TFE was still that preferred for the PCL-based solution. In the “supplementary data section”, details of the electrospinning parameters for each scaffold, based on size and type of polymers, are presented as a table.

#### 2.1.2 Scaffold surface treatment: Plasma functionalization and plasma polymerization

It is known for many years that synthetic polymers are characterized by low surface energy and low wettability, leading to weak adhesion with other surfaces, including living cells. This can be remedied by modifying the polymer surface using exposure to low-temperature (non-equilibrium) plasma [45]. In this present work, cell adhesion to the hydrophobic surfaces of pristine electrospun scaffolds is generally very limited [44]. To correct this shortcoming, the surface compositions of scaffolds were modified by incorporating selected new functional groups using plasma treatment with different gases or gas mixtures. Treatment or coating uniformity throughout the (typically 250 μm-thick) 90% porous scaffolds are assured by the fact that its open volume facilitates diffusive transport of active precursor species from the plasma. This fact and details regarding plasma apparatus and methodology have been validated and presented elsewhere[44, 46-49]. We have also shown previously that plasma activation of electrospun scaffolds greatly enhances cellular interactions with those scaffolds that are highly favourable and conducive to tissue engineering applications [44, 47].

Here, the surfaces of the fibers were exposed to low-pressure radio-frequency (rf) glow discharges in selected gases and also coated with an ultra-thin (< 100 nm) amine-rich plasma-polymer films symbolized by “L-PPE:N”. Since the details have been described in previous work, we explain here only the most essential aspects. The surface of nanofiber mats was functionalized by using oxygen or ammonia gas (O_2_ or NH_3_, both from Air Liquide Canada Ltd., Montreal, QC) in a low-pressure (600 millitorr or 80 Pa) capacitively coupled radio-frequency (r.f., 13.56 MHz) glow discharge plasma reactor (cylindrical aluminum/steel chamber) with a gas flow rate of 15 standard cubic centimeters per minute (sccm) and plasma exposure time of 30 s for O_2_ and 1 min for NH_3_ under mild plasma condition (power: *P*=15 W and self-bias voltage: *V*_b_=-40 V). Alternatively, plasma-polymerized “L-PPE:N” coating was deposited on the scaffold surface in the same low pressure (80 Pa) glow discharge plasma reactor by using ethylene (C_2_H_4_)/ammonia (NH_3_) gas mixture (C_2_H_4_: 99.5%; NH_3_: 99.99%), also under mild plasma conditions (*P*: 15 W, *V*_b_: -40 V). The flow rates of the two high-purity feed gases, C_2_H_4_ and NH_3_, were kept at 20 sccm and 15 sccm, respectively for a duration of 7.5 min for both sides of the scaffold samples. These conditions had been previously optimized to yield a thin layer of almost 100 nm of L-PPE:N on a glass microscope slide, with adequate nitrogen ([N]) and amine concentrations [NH_2_] on the surface, and low solubility in cell culture media [48, 50]. The treated samples were finally sealed in sterilized Petri-dishes for the subsequent steps.

### 2.2. Scaffolds Characterization

#### 2.2.1. Scanning Electron Microscopy (SEM)

To evaluate electrospun nanofiber structure and morphology, the surface of polymeric mats was characterized by Scanning Electron Microscopy (SEM) using a Hitachi model TM3030plus instrument at a working distance of 2 mm and voltage of 15 kV, performed at different magnifications. Selected samples were mounted on a sample holder using double-sided adhesive tape and adjusted at the targeted distance. The diameters of 200 randomly selected fibers for each sample of large, medium and small fiber size (at least three different spots/sample in triplicate) were measured either *in situ*, or captured SE-micrographs were then analyzed using ImageJ analysis software.

#### 2.2.2. Overall porosity and pore size distribution

Overall porosity of PLA electrospun mats of the three different fiber sizes (small, medium and large) was quantified using a liquid (ethanol) intrusion method [51, 52]: Dry mats were first weighed (W1) before being immersed in 100% ethanol overnight for complete wetting of the samples. Next day, the mats were gently wiped to remove excess ethanol and weighed again (W2). Porosity is defined as the volume of ethanol trapped in the mats (∼[W2-W1]) divided by that of the wet mats (∼W2). This technique, as well as mercury intrusion porosimetry, had been verified in previous studies [44, 46] and both had been found to yield very similar results. In addition, calculation of average pore size and its distribution in the nanofiber scaffolds for each fiber size, was performed by using an useful and practical plugin named “DiameterJ” in ImageJ software, through fitting an ellipse inside the pores’ shape, and the mean pore size for each scaffold (from three different spots per sample in triplicates) was measured from the average of the long and the short axes for each fitted ellipse.

#### 2.2.3. Mat thickness measurement

The thicknesses of nanofiber mats (targeted design values at 250 μm) were determined using a digital micrometer gauge for film thickness measurements (Film Master, Qualitest, 10 μm resolution). The samples were sandwiched between two rigid PET films to minimize errors resulting from compression during measurements.

#### 2.2.4. Surface chemical analyses (XPS)

X-Ray photoelectron spectroscopy (XPS) analysis was performed on treated and non-treated scaffolds in a VG ESCALAB 3MkII instrument, using non-monochromatic Mg Kα radiation. The sampling depth, in the range of 1-5 nm, depends on the fiber geometries in the ca. 1mm^2^ analyzed area. To acquire spectra, emission angle was set at 0^0^, normal to the mat surface, and charging was corrected by referencing to the C1s peak at binding energy BE=285.0 eV. The X-ray source’s operation condition was at 15 kV, 20 mA and quantification of the constituent elements was performed using Avantage software (Thermo Electron Corporation) after Shirley-type background subtraction, following which the concentrations of elements were determined from XPS survey spectra.

#### 2.2.5. Static contact angle measurements

Static contact angles (SCA) of water droplets were measured at room temperature using a FDS tensiometer, OCA Data Physics, model TBU 90E. Treated and non-treated mat samples were fixed on glass slides; then 2μL droplets of MilliQ water were placed on the sample surfaces with a micro-syringe (at least four different spots/sample, carried out in triplicate) and average values along with standard deviations of SCAs were evaluated using SCA20-U software provided by the manufacturer.

### 2.3. Biological Experiments

#### 2.3.1. Cell culture, seeding and preparation of the 3D models

The epithelial breast cancer cell line MDA-MB231 (Green-fluorescent protein, GFP), and stromal connective tissue cells including IMR-90 mcherry Fibroblasts (Red-florescent protein, RFP) [53, 54], provided by the laboratories of professor M. Park at McGill university, were cultured in high-glucose Dulbecco’s modified Eagle medium (DMEM), supplemented with 10% fetal bovine serum (FBS), 1% Pen Strep (PS) antibiotic (all from Gibco, Thermofisher), at 37 ^0^C in a humidified cell culture incubator with an atmosphere of 5% CO_2_. After reaching 90% of confluency, cells (passage 3) were first washed with sterile phosphate buffered saline (PBS), detached using 0.25% trypsin (Gibco, Thermofisher), followed by adding fresh RPMI cell culture medium (Gibco, Thermofisher) supplemented with 10% FBS and 1% PS. Then, the cell suspensions were centrifuged at 1500 rpm for 5 min to collect cells at the bottom of vials, and finally suspended in 3 mL fresh complete media, counted and diluted to 100,000 cells/mL for fibroblasts, ready to be seeded on the scaffolds and 500,000 cells/mL for cancer cells to be mixed with hydrogel. Treated and non-treated electrospun mat scaffolds were precisely cut into disks with a 9 mm punch, sterilized with RPMI media containing 1% antibiotic to remove possible contaminants, then finally placed into individual wells of non-adherent 48-well polystyrene culture plates, destined here for i) a tumor migration assay; ii) a high-throughput therapeutic screening test. Next, 200 μL of fibroblast cell suspension (20,000 cells/scaffold) was added to each well and incubated for 30 min in the humidified atmosphere of 5% CO_2_. Then, medium was aspirated from each well and rinsed with fresh media to remove non-adherent cells. Finally, a layer of (100 μL) of alginate / gelatin hydrogel (A1G7), pre-impregnated with MDA-MB 231 tumor cells (50,000 cells in gel/well), was applied on top of each sample compartment, followed by adding 200 μL of CaCl_2_ for ionic crosslinking. Each well was then aspirated, washed twice with fresh media to remove residual crosslinking agent and 500 μL of complete media was finally added per well. The plates were incubated for 1, 3 and 7 days, the culture medium being changed every three days.

#### 2.3.2. Investigation of Initial Cell adhesion

Initial adhesion of fibroblasts seeded on the treated and non-treated scaffolds with different pore and fiber sizes was assessed after 30 min incubation of the cells. After this, the fibroblasts-seeded scaffolds were fixed in 4% paraformaldehyde solution (PFA) diluted in DI water for 15 min at room temperature, before washing twice with PBS solution. They were subsequently placed on glass slides for observation using an EVOS M5000 fluorescence microscope. Images of the whole surface of each 9 mm-disk were captured with a 4X-objective.

#### 2.3.3. Tumor cell migration - observation and quantification

Movement and migration of cancer cells was assessed as a function of incubation time. The top surface and depth of treated and untreated scaffolds (nominal thickness 250 μm) were monitored after 1, 3 and 7 days, after scraping off the hydrogel and fixing the surface with 4% PFA. The samples had been placed onto microscope slides, first covered with a droplet of mounting medium (containing DAPI) to avoid dehydration, then with a protective glass cover slip. Images of green (tumor cells) and red (fibroblast) cells on the top surface and within the scaffolds were captured under different magnifications using a florescent and a confocal scanning microscope, namely EVOS M5000 (4X) and Olympus IX81 (10X), respectively. Images were analyzed and the florescent intensity and/or the number of migrated cells counted and quantified using ImageJ software. Each experiment was done in triplicate, with quantification from 20 spots to fully cover the surface of each sample.

#### 2.3.4. Drug screening experiment

A drug screening experiment was devised using a well-known cancer drug, Doxorubicin (Sigma), to evaluate tumor migration in our 3D cell culture model. 48-well plates loaded with O_2_ plasma-treated PLA mats (medium size), as before pre-cultured with Fibroblasts and tumor cells, were treated either (i) with various concentrations of Doxorubicin, namely 0, 0.05, 0.1, 0.5, 1 and 2 μM; or (ii) with sterile PBS as control, in low-serum conditions (1% FBS) in triplicate for 7 days. The media loaded with the drug was replaced on day 4 for each well. An alternative drug screening experiment was also carried out, namely “Boyden Chamber Migration Assay” [20, 21]. based on Matrigel^®^, with the aim of comparing the results with our present 3D model. Similar numbers of fibroblasts (20,000 cells/well) were seeded in a 24-well plate and 500 μL of complete media (10% FBS) were added to each well. In the second 24-well plate, Matrigel^®^ with MDA-MB 231 breast cancer cells (50,000 cells/well), in triplicate, was loaded in the upper compartments of Boyden trans-well chambers with suitable transparent PET membranes (8 μm pore size; Corning, NY, USA); following this, 500 μL of complete media (10% FBS) was added and incubated at 37 °C, 5% CO_2_. After 24 hours of incubation, PET membranes with Matrigel^®^ and tumor cells were transferred to the first plate with the fibroblasts, and cells were then treated with combined PBS / Doxorubicin at concentrations of 0.05, 0.1, 0.5μM in RPMI media with low-serum conditions (1% FBS) in the upper compartment. Tumor cell migration was triggered over 7 days using RPMI supplemented with 2% FBS as a chemoattractant in the lower compartment.

### 2.4. Statistical Analysis

All experiments reported here were carried out in triplicate to evaluate reproducibility and all data were expressed as mean values ± SD/SE. Statistical analysis was carried out using two (or three)-way ANOVA with Tukey’s post hoc analysis for parametric data; in the case of having non-parametric data Kruskal-Wallis test was done, followed by Mann-Whitney post hoc analysis to compare two independent groups of interest. Values of p < 0.05 were considered significant for all tests.

## 3. Results and Discussion

As illustrated in the schematic diagram, Figure 1, *PP-3D-S* fabrication involves several steps, namely (i) electrospinning of a suitable biodegradable polymer (herein mainly PLA) as scaffold; (ii) bio-activation via treatment of the nanofiber scaffold by plasma functionalization or ultra-thin plasma polymer (PP) coating; (iii) seeding stromal cells within the open 3D volume of treated scaffolds to mimic target tissues; and finally (iv) depositing a droplet of hydrogel pre-seeded with different types of tumor cells on top, to evaluate tumor cells migration and metastasis over specific time intervals. Results of experiments and discussion are presented in the following text.

### 3.1. Physical properties and morphology evaluation of nanofibrous mats

#### 3.1.1. Fiber diameter and distribution

Electrospun (PLA) scaffolds have been used frequently in bone tissue engineering [55] and 3D cancer modeling [56]. These can be made from PLA, but also from other polymers like polyurethane (PU) or poly-caprolactone (PCL); they can be fabricated with different morphological features including pore size, porosity, and fiber diameters, as well as various mechanical properties, mimicking different target tissue matrices. We have previously developed these scaffolds for neuronal [57], cardiac [58], vascular [44, 47] tissue engineering applications, but now are developing this toward modelling a tumor microenvironment. Figure 2 shows micrographs of electrospun mats captured by SEM, along with the plots of fiber diameter distribution and the average size (three individual experiments with triplicate samples), in which the red dotted line presents the average (at peak). The micrographs clearly demonstrate randomly interconnected network structures of the electrospun mats, which proves the desired smooth surfaces lacking “beads” (spheroidal-shaped defects), and thus witnessing favorable processing conditions. As targeted, the value of fiber diameter for PLA mats was mainly found to be in the range of 200-300 nm for “Small” (average diameter: 273±30 nm), 600-800 nm for “Medium” (average diameter: 730±75 nm), and 1-2 μm for “Large” (average diameter: 1.47±0.2 μm) scaffold size; for the cases of PU and PCL mats, the mean values for medium size were 820±40 nm and 910±86 nm, respectively. SEM characterization was also done on plasma-treated mats, and as expected (images not shown) no perceptible difference was observed between bare (non-treated) and treated ones, as also reported in previous work[44].

**Figure 2.**
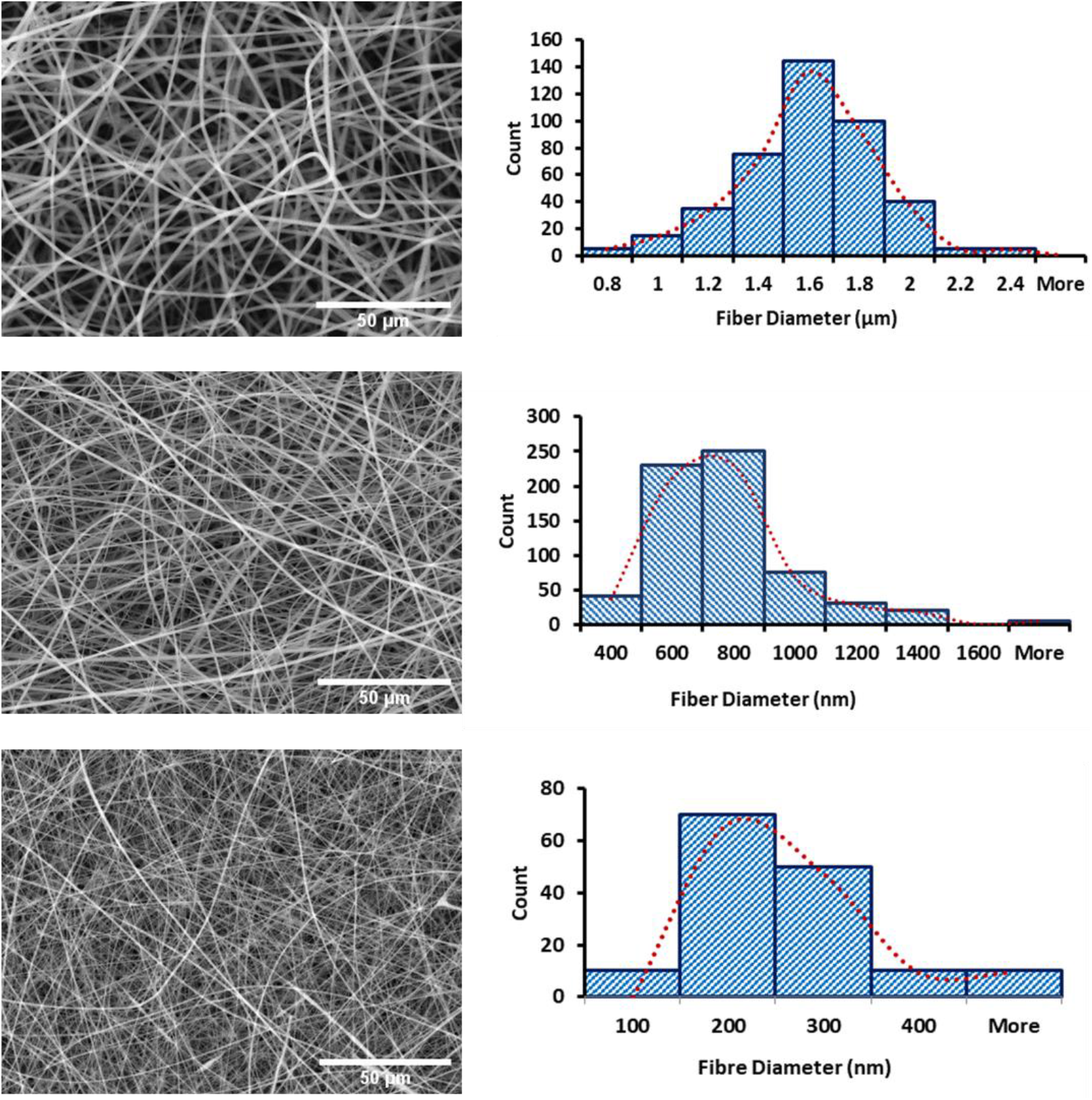
SEM micrographs and fiber diameter distribution curves of PLA electrospun mats of different fiber diameter sizes (small:200-400 nm, medium:600-800 nm, large:1-2 μm), scale bar:100 μm

#### 3.1.2. Thickness of nanofibrous scaffolds

The final product is an electrospun sheet with dimensions of 28 by 30 cm in length and width (mandrel diameter: 10 cm), respectively. Thickness measurements, using a digital micrometer, usually vary substantially from the center (max. 280 μm) to the edges (ca. 80 μm). Hence, samples (squares of 5 by 7 cm, 200-250 μm) were taken from central portions of the sheets for use in the subsequent experiments.

#### 3.1.3. Porosity and pore size measurement

Table 1 and Figure 3 show overall porosity and average pore sizes of PLA mats (small, medium, and large sizes) obtained by the methods in section 2.2.2 (liquid intrusion technique and ImageJ). The so-called “large” scaffolds possess larger pores and slightly smaller overall porosity than the other two; the observed high porosity of ca. 90% in all three cases is of course highly desirable for intended use, because it readily enables transport of oxygen and other gases, nutrients, metabolic wastes, etc. during metabolic activity in the biological microenvironment. Furthermore, the (multi-μm) pore sizes are clearly large enough to allow ready accommodation of stromal cells into the scaffolds. To verify data obtained by the two above-mentioned methods, we also resorted to others, including MIP (Mercury Intrusion Porosimetry) test; porosity (ε) was evaluated gravimetrically according to equation (1) [59], and average pore diameter (r) by equation (2)[60].

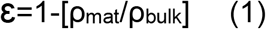

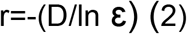

**Table 1.** Comparison of average pore size (μm) and porosity (%) of PLA electrospun mats in different sizes calculated by three techniques including ImageJ, eq. (2) and MIP test

| Parameter | Scaffold | ImageJ | eq. (2) | MIP Test (30Kpsia) |
| --- | --- | --- | --- | --- |
| Average Pore Diameter, $r$ ($\mu\text{m}$) | Large | 3.0 ± 0.32 | 5.2 ± 0.6 | 2.94 |
| | Medium | 2.1 ± 0.2 | 3.3 ± 0.5 | 2.91 ± 0.6 |
| | Small | 1.0 ± 0.05 | 1.3 ± 0.1 | 1.99 |
| Porosity (%) | Large | 90.3 ± 1.5 | 87.4 ± 0.45 | 84.2 |
| | Medium | 91.0 ± 1.6 | 87.2 ± 0.3 | 84.0 ± 1.05 |
| | Small | 91.3 ± 0.9 | 88.3 ± 0.25 | 86.1 |

**Figure 3.**
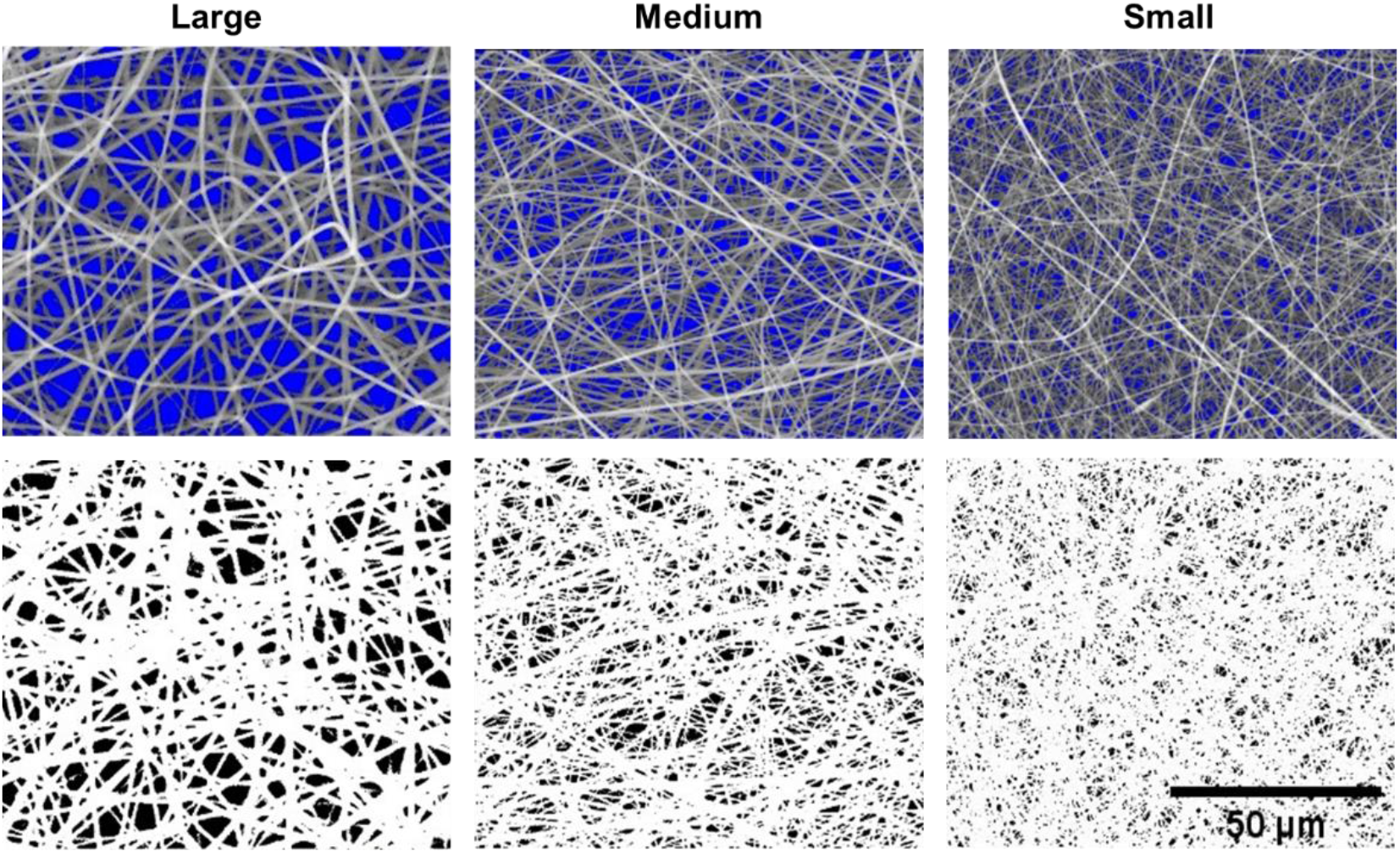
SEM micrographs of electrospun PLA mats with different fiber sizes and porosities (large, medium and small fibers), scale bar:100 μm

where ρ_mat_ is the electrospun mat’s measured density, ρ_bulk_ is the bulk density of PLA (1.24 g/cm^3^), and D the mean fiber diameter. As shown in Table 1, there are slight differences between average pore diameters obtained from ImageJ and eq. (2). Pore diameters for medium and large mats determined by the MIP test were found to differ little, possibly for the following reason: It seems that pores in the bulk differ in size from voids observed from SEM images. Possibly, during mercury intrusion into the mats, the latter can deform and enlarge the apparent pore size. Nevertheless, data from Figure 3 and Table 1 confirm that the present PLA mats are suitable 3D networks for accommodation and displacement of live cells through the nanofibrous structures, as they do in their natural ECM, both the stromal fibroblasts as well as MDA-MB 231 breast cancer cells, respectively 10-15 and 15-20 μm in average size [61, 62].

### 3.2. Surface characterization of bare and plasma-treated scaffolds

#### 3.2.1. Surface-Chemical Analyses (XPS)

In section 2.1.2 we have already described the rationale for various plasma treatments used in the current context of 3D nanofibrous scaffolds. Following treatments, XPS analyses were used to evaluate surface chemical compositions of the substrates that had been treated with O_2_ plasma or by L-PPE: N coating, alongside pristine PLA mats. XPS survey spectra (not shown) indicate surface-near heteroatom concentrations before and after treatments, as presented in Table 2. As expected, the results in this table show increases in the [O] and [N] concentrations of scaffolds treated with O_2_ and L-PPE:N, respectively, in comparison with pure PLA mat, confirming the presence of oxygen- and/or nitrogen (amine)-rich functionalities on the surface of scaffolds. Detected fluorine (F) originated from the Teflon-sprayed aluminum foil used for collecting nanofiber mats while electrospinning; some bound F, like here, is considered to be safe, without danger of cytotoxicity and of no risk to cells during culture.

**Table 2.** Surface chemical compositions of pure PLA and PLA mats modifieded with L-PPE:N coating or O_2_ plasma, determined from XPS survey spectra

| ELEMENT | Pure PLA (At. %) | O<sub>2</sub> Plasma (At. %) | L-PPE:N Coating (At. %) |
| --- | --- | --- | --- |
| F1s* | 13.3 | 13.2 | 2.2 |
| O1s | 36.6 | 39.8 | 4 |
| N1s | - | - | 12.7 |
| C1s | 50.1 | 47 | 81.2 |
| [O]/[C] | 0.73 | 0.85 | 0.05 |
| [N]/[C] | - | - | 0.16 |

\* The source of fluorine is the Teflon spray applied to the Aluminium foil used to collect nanofibers during the electrospinning process.

#### 3.2.2. Static Contact Angle (SCA) Measurements

SCA results are summarized in Table 3. Although these must be considered only as qualitative in view of the porous, rough surface, it is obvious that medium PLA samples treated with O_2_ and NH_3_ plasma dramatically change surface properties from hydrophobic to fully hydrophilic (0°) in comparison with the pristine PLA mat: Immediately upon contact, water droplets spread and diffused into the plasma-treated mats. Capillarity, surface roughness, porosity, and high polarity due to chemically-bonded oxygen (Table 2) all combined to cause the observed increase in wettability, but surprisingly this was not the case for samples coated with L-PPE:N, which is known to include desired bioactive nitrogen-rich functionalities such as primary amines.

**Table 3.** Static water contact angles of plasma-treated and non-treated PLA mats

| Samples | Static Contact Angle (°) |
| --- | --- |
| Pure PLA | 142 ± 2 |
| O<sub>2</sub> | 0 |
| NH<sub>3</sub> | 0 |
| L-PPE:N | 132 ± 3 |

### 3.3. Investigation of initial cell adhesion

#### 3.3.1. Effect of treatment type and scaffold size on stromal fibroblast adhesion

First, the effects of three different plasma treatments and three mesh sizes were studied with regard primarily to bonding of cancer-associated stromal fibroblasts, for both treated and non-treated PLA mats, after 30 minutes of incubation. Because pristine PLA mats are hydrophobic, they do not facilitate cell seeding and adhesion, unlike all plasma-treated scaffolds, as illustrated in Figure 4. Statistically significant differences are noted between the percentages of cells adhered to the scaffolds with different morphologies and plasma treatments (H=27.33, P=0.0041). The observed greatly-improved cell adhesion on plasma-modified scaffolds (NH_3_ and O_2_, compared with Ctrl) can be respectively attributed to positively- or negatively-charged hydrophilic functional groups [47]. Indeed, we had already much earlier reported that cell adhesion and -proliferation on plasma-modified electro-spun 3D scaffolds greatly exceeded those on virgin control samples [44, 46]. The results shown in Figure 4 present *surface-near* numbers of the cancer-associated fibroblasts, because merely 30 minutes after seeding no appreciable cell penetration deeper into the porous scaffold should be expected for small-, medium, or large-sized fiber scaffolds, plasma-modified or not. Interestingly, although L-PPE:N-coated samples displayed unexpected “hydrophobic” behaviour (see Table 3), their bioactive amine groups nevertheless are seen to have promoted fibroblast adhesion, albeit less pronounced than for the O_2_ and NH_3_ plasma-modified cases. Finally, the medium and large-sized scaffolds displayed much higher cell counts than their small-sized counterparts, presumably owing to much larger pore sizes even near the surface (see Table 1).

**Figure 4.**
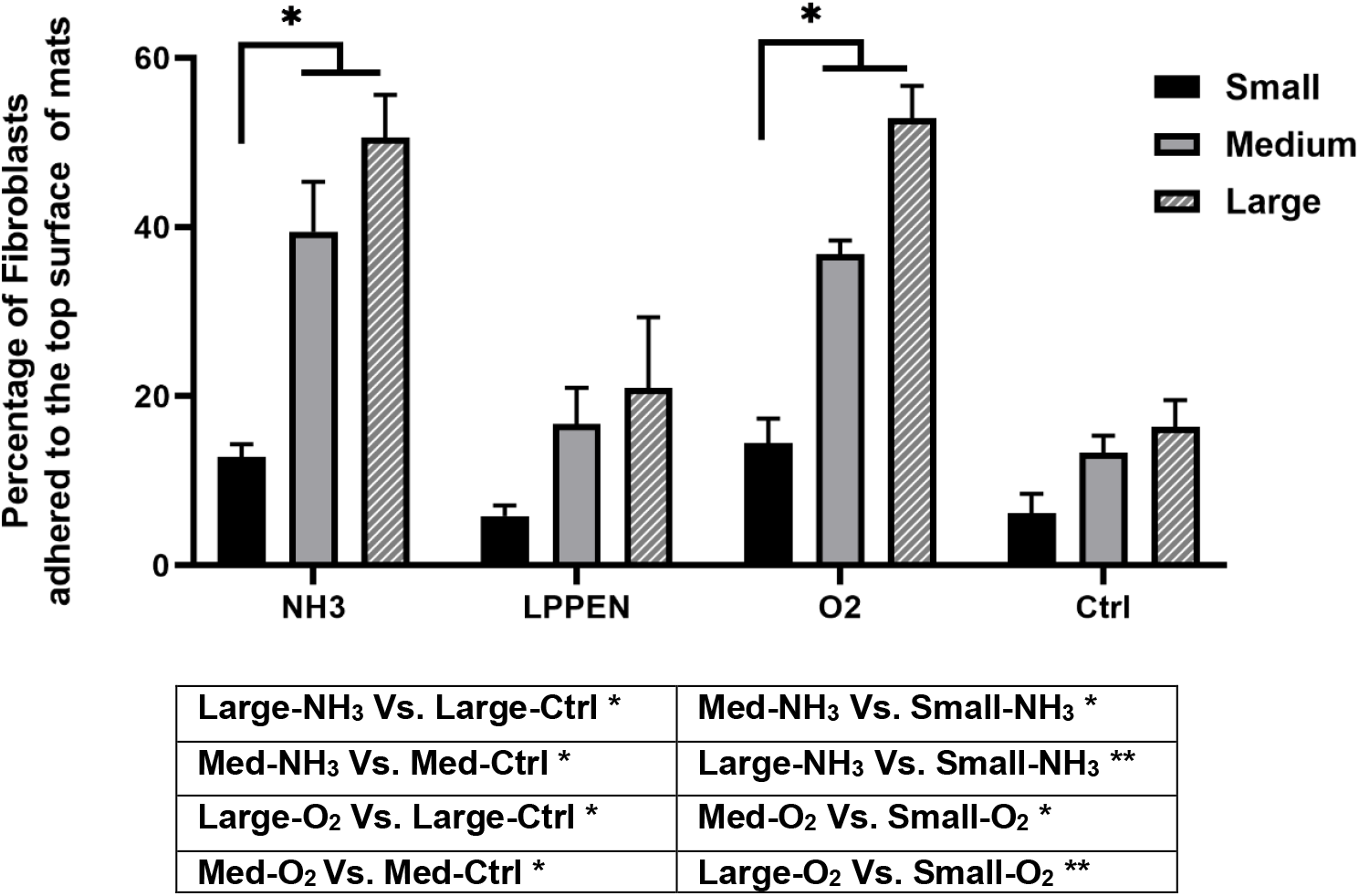
Percentages of initially-seeded RFP cancer-associated fibroblasts adhering to plasma (O_2_, NH_3_ and L-PPE:N)-treated and non-treated PLA scaffolds 30 minutes after seeding 20,000 cells (error bars: SE., n=3)

### 3.4. Tumor-cell migration

#### 3.4.1. Effect of plasma treatment type on tumor cell migration

We next investigated migration of MDA-MB-231 breast cancer cells into the 3D microenvironment of our PP-3D-S model. Breast cancer commonly metastasizes to other organs or to bone, for example to the spine [63], so we have conducted a series of experiments that elegantly simulate breast cancer metastasis. Over 7 days, the GFP triple-negative breast cancer cells, seeded within the upper hydrogel layer, readily migrate to the medium-sized PLA mat first seeded with IMR-90 mCherry fibroblasts (Figure 5). Figure 5(A) presents microscopic images of invading and migrating (and proliferating) breast cancer cells (green) in the 3D scaffolds, in which the RFP fibroblasts (red) were pre-adhered to different plasma-treated (O_2_, NH_3_, L-PPE:N) and untreated medium-sized PLA scaffolds. The reader is reminded that the tumor cell-bearing hydrogel layer was removed by scraping just prior to acquisition of the images; therefore, these micrographs manifest greatly increasing presence of the (green) tumor cells well within the scaffold’s volume, by having changed the type of treatment and also by increasing the incubation time (statistically proved: H(12)=33.4, P=0.00045). This is clearly observable in all four cases (including untreated controls), especially after 7 days, as also quantitatively illustrated in the lower portion, Figure 5(B). Now, referring to Figure 5(A), O_2_ and NH_3_ plasma-treated scaffolds very evidently show quite large (green) tumor cell populations after 7 days compared with untreated controls, especially NH_3_ (O_2_ displaying quite significant scatter among the n=3 data sets). But the unexpected, surprising result is that relating to the L-PPE:N coated scaffolds, which show only marginal improvement over corresponding untreated control samples. We shall return to this perplexing observation in the Discussion section later in this text.

**Figure 5.**
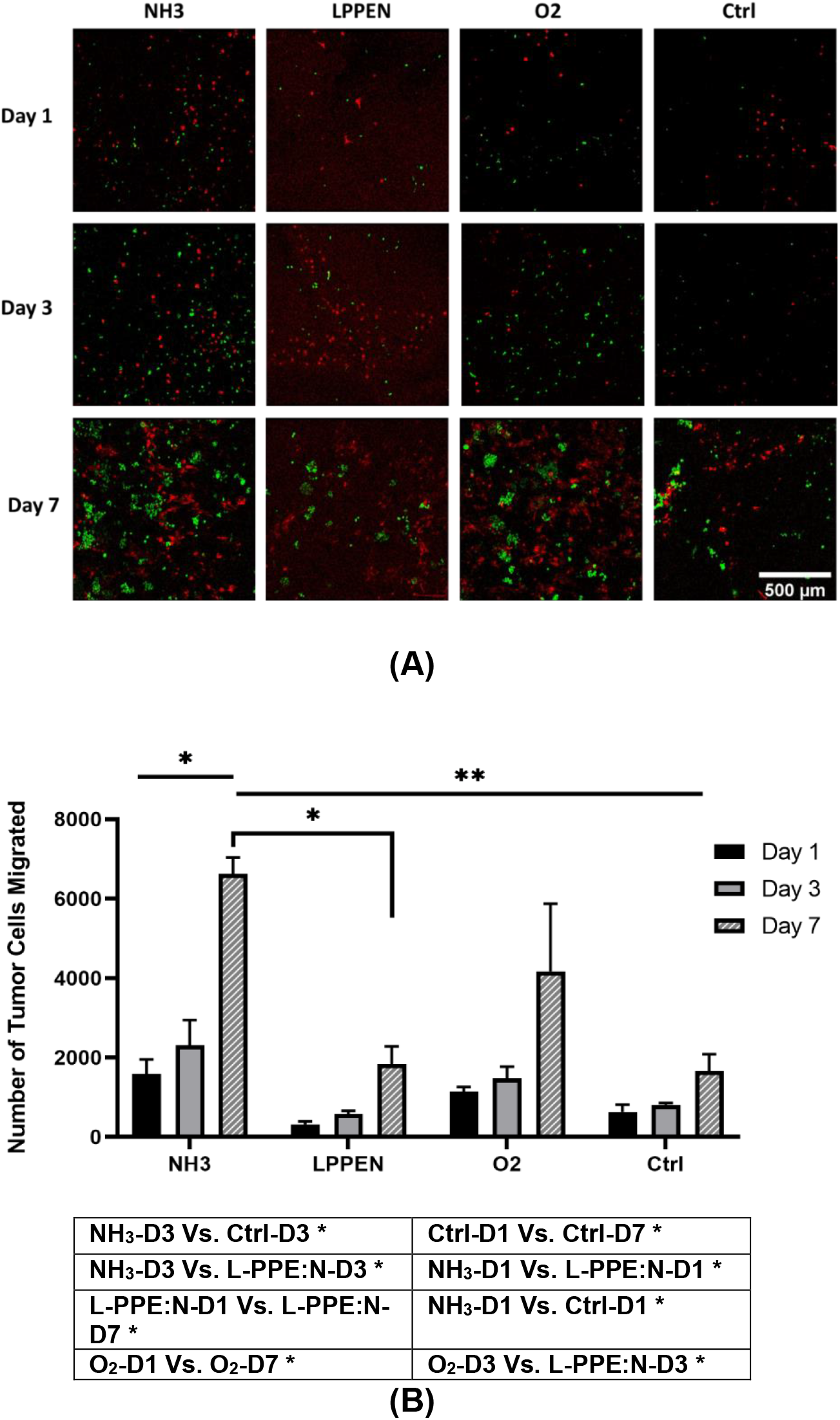
Tumor cell migration in the PP-3D-S model. (A): Representative images of RFP cancer-associated fibroblasts (red) adhering to untreated controls, and to L-PPE:N, O_2_, NH_3_ plasma-treated medium-sized PLA fiber scaffolds, along with increasing numbers of GFP breast cancer cells (green) having migrated over periods of up to 7 days. (B): histograms that quantitatively represent tumor cell migration following the different types of plasma treatments and durations (error bars: SE., n=3)

#### 3.4.2. Effect of scaffold size on tumor cell migration

Figure 6 shows not only the effects of plasma treatment types on tumor cell migrations after different incubation periods (see Figure 5(A,B)), but here also that of the three different scaffold sizes, small, medium and large. Many of the comments made in preceding section 3.4.1. for the case of the medium-sized scaffolds are seen to apply here, too, namely large numbers of migrated / proliferated (green) tumor cells observed in the scaffold volumes, especially after 7 days for O_2_ and NH_3_ plasma treatments, while L-PPE:N results only marginally improved compared with their control counterparts. The principal new observations (statistically significant difference: H(35)=96.95, P<0.00001) that may be noted in Figure 6 are: i) Small-sized scaffolds, even those corresponding to O_2_ and NH_3_ plasma treatments, display only very minor variations among themselves, compared with untreated control samples; ii) Large- and medium-sized scaffolds show rather similar numerical values, slightly higher for “large” in the NH_3_ and L-PPE:N cases but not significantly different compared with “medium”. Data from Figure 6 indicates that the significantly reduced pore openings that characterize small-sized scaffolds (ca. 1 μm, compared with roughly 2 and 3 μm for medium- and large-size, respectively, see Table 1) evidently greatly hindered cell mobility through the open volumes of the “small” scaffolds, even though those free volumes were comparable in all three cases, roughly 90% (Table 1). Not even the much smaller fiber diameters (Figure 2) and correspondingly greater flexibility of the small-sized scaffolds’ fibers could compensate for obstacles posed by the reduced pore volumes. This observation somewhat parallels that of fibroblast adhesion on the topmost surface layer of the three different-sized scaffolds (see section 3.3.1., Figure 4), where small scaffold size initially (first 30 minutes) led to much lesser numbers of adhering cells in all cases.

**Figure 6.**
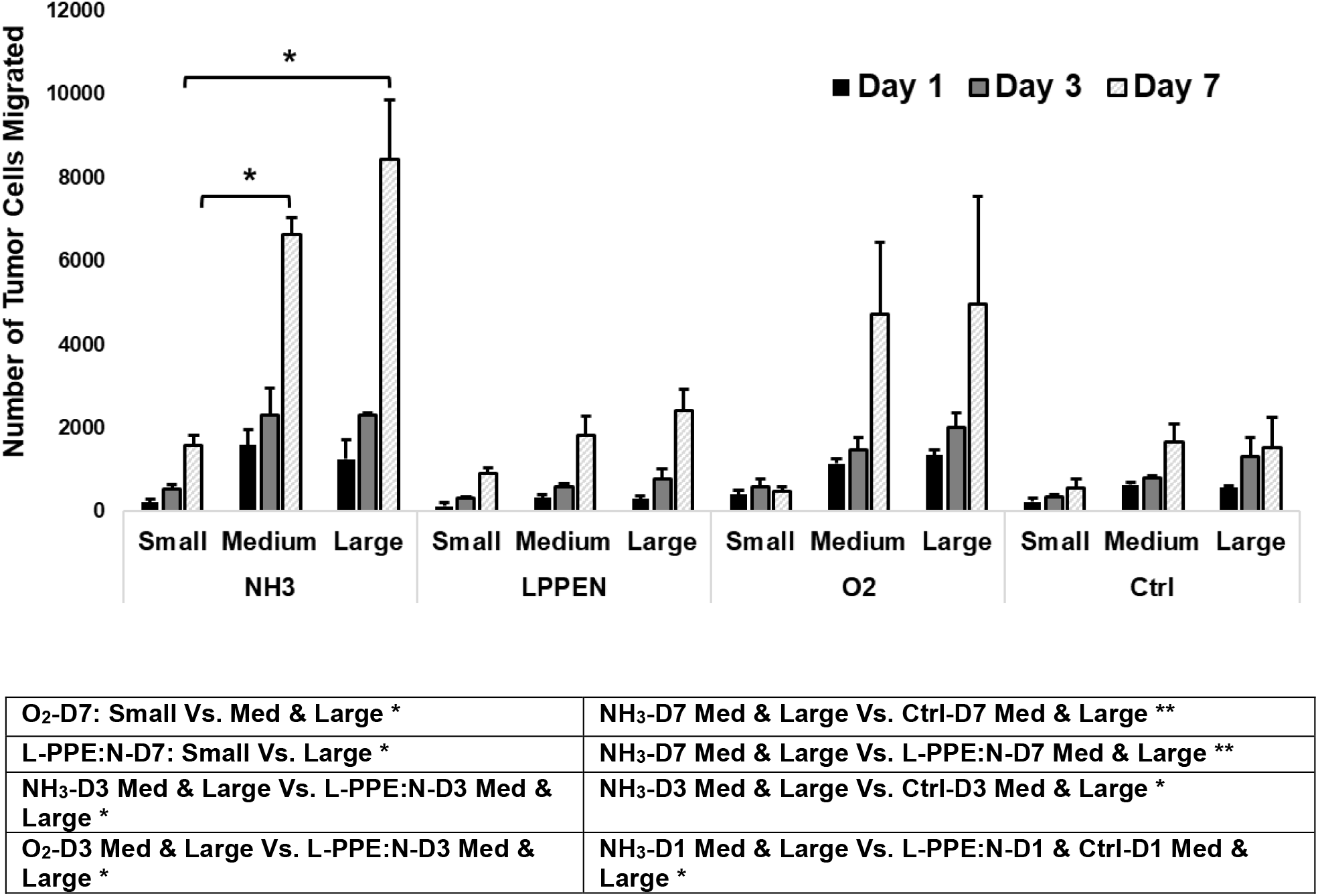
Numbers of tumor cells having migrated into the volume of different-sized and differently plasma-treated scaffolds (error bars: SE., n=3)

#### 3.4.3. Effect of polymer type on tumor cell migration

As pointed out in section 2.1.1., electrospun mats of two other polymers besides PLA were also fabricated, namely polyurethane (PU) and poly(caprolactone) PCL, all of “medium” size (fiber diameters in the 800 – 900 nm range, and all having here undergone identical O_2_ plasma treatments). Those three materials possess among themselves quite different mechanical stiffness characteristics. In Figure 7 are presented (A) micrographs that show the red- and green-fluorescent fibroblasts and tumor cells, respectively; and (B) the corresponding numbers of tumor cells found migrated / proliferated in the scaffold volumes after 1, 3 and 7 days. What most clearly emerges here is that after 7 days, no statistically significant differences (F(2,1.7)=0.834, P=0.5593; W(2,1.5)=1.165, P=0.4950) can be observed among the three cases, in spite of the appreciably differing fiber stiffnesses. Comparing this outcome to that of preceding section 3.4.2, one may conclude that the main parameter which clearly dominated the tumor cells’ mobilities / proliferations, hence their numbers within the scaffold interiors, was the scaffold morphology; by this is meant the combination of average pore and fiber sizes, free volumes and numbers of O-based surface-functional groups from plasma treatments presumably being very similar. In other words, mechanical stiffnesses of the electrospun polymers somewhat surprisingly appear to have played only a relatively minor role. This demonstrates that the present PP-3D-S tissue model can potentially be readily fabricated based on different (biodegradable) polymers having different mechanical and/or other physical properties, tuned to best match the targeted tissue types (bone, soft tissue, ⃛).

**Figure 7.**
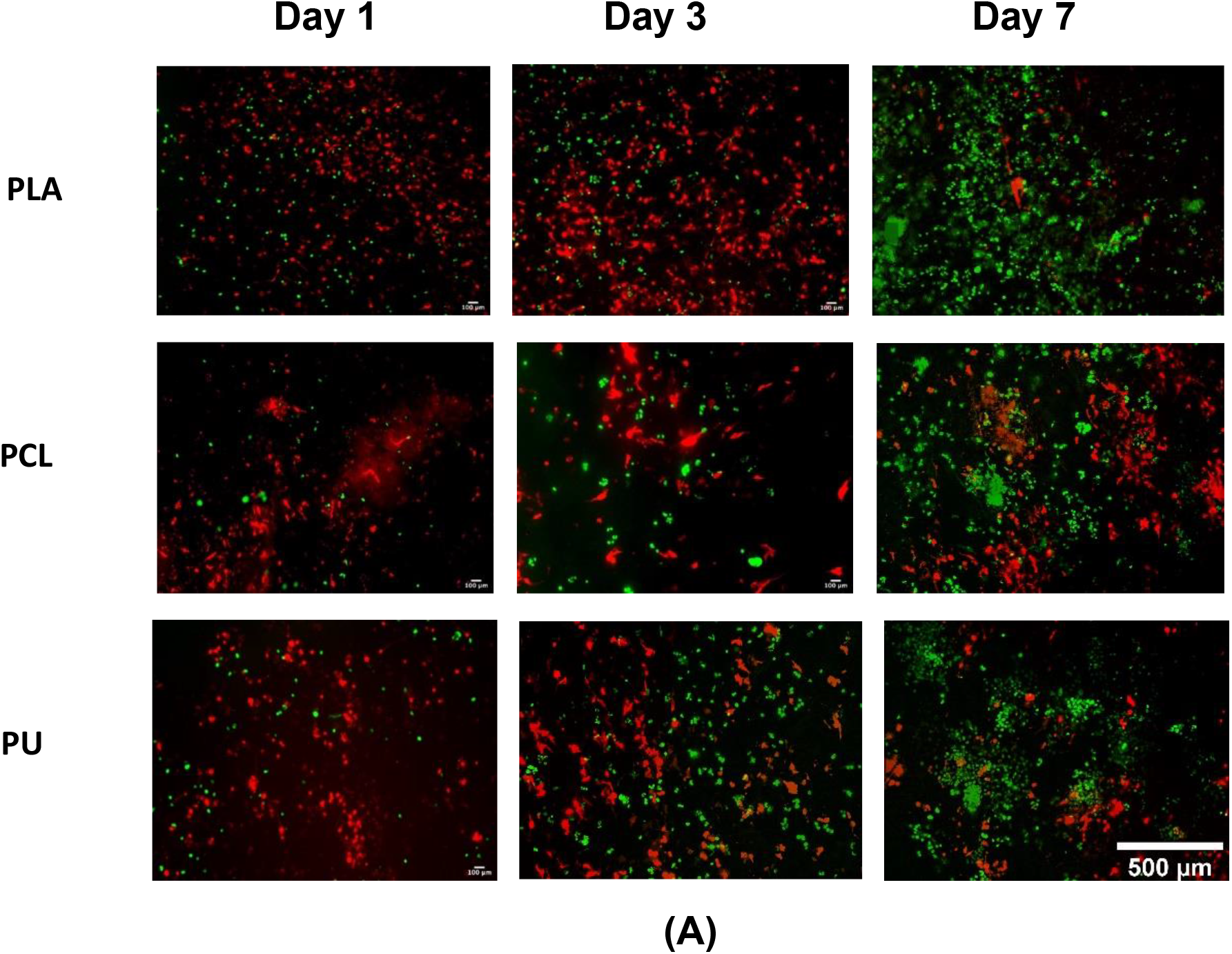
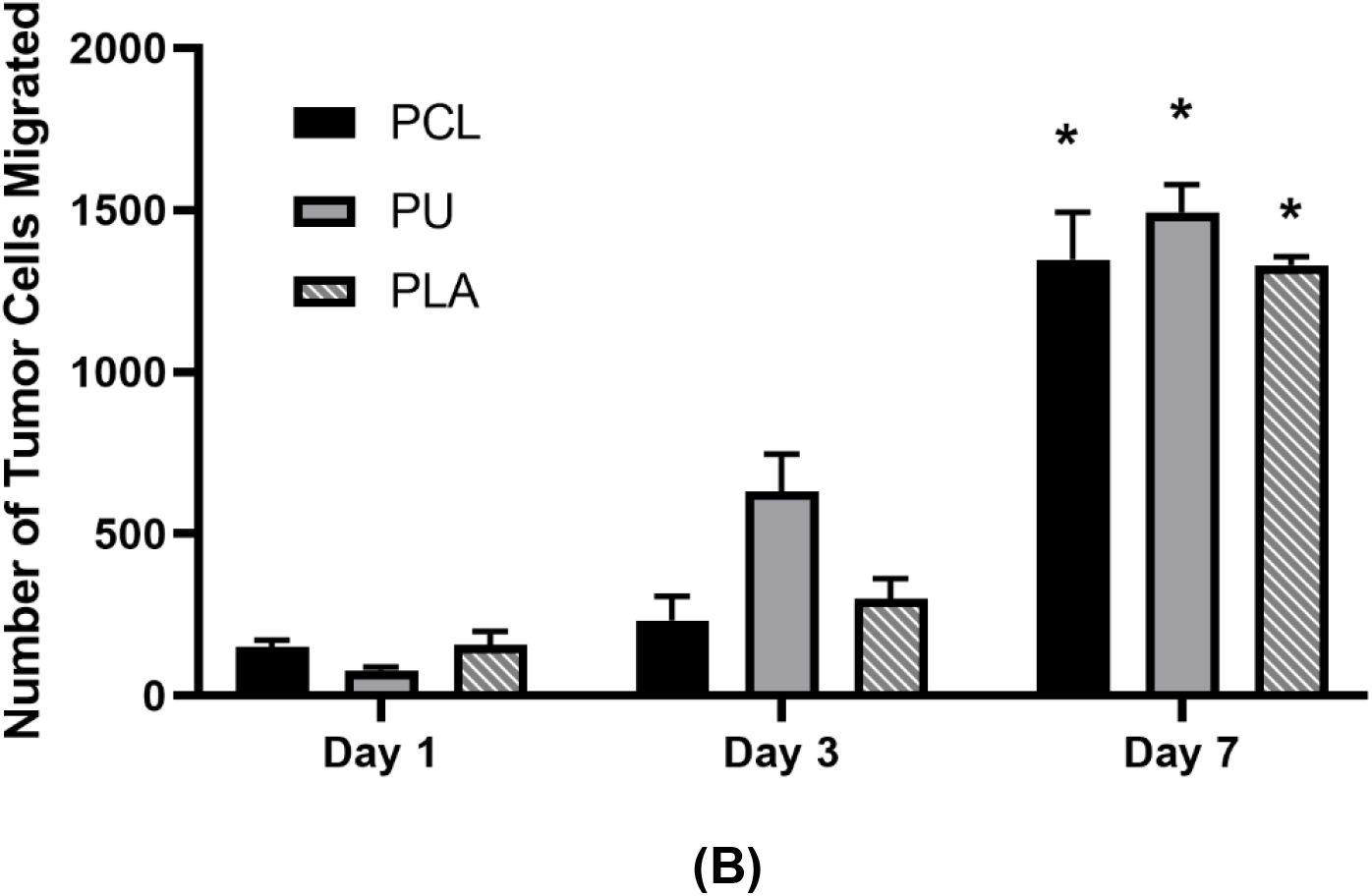
(A) optical micrographs showing (red) fibroblasts and (green) tumor cells; (B) Tumor-cell numbers after 7 days of migration and proliferation in the volumes of “medium”-sized scaffolds of three different polymers, all O_2_ plasma-treated (**The only pairs with P<0.05:** PCL: D1 Vs. D7; PU : D1 Vs. D7; PLA : D1 Vs. D7; error bars: SE., n=2)

### 3.5. Penetration depth of tumor cells

Top-view photomicrographs of red- and green-fluorescent cells shown in preceding sections give no indication how deep below the top surface of the ca. 250 μm-thick 3D scaffold those cells were located. However, that information can be derived from confocal microscopic observations portrayed in Figure 8(A), where a 3-dimensional distribution is shown that also includes the z-direction (blue arrow), down to a maximum depth of ca. 120 μm below the original hydrogel-mat interface (z=0), for all plasma-treated and non-treated medium-sized scaffolds after 7 days. In Figure 8(B) the tumor cell migration into the depth of NH_3_- and O_2_ plasma-treated mats was significantly greater than that observed for L-PPE:N-coated and -untreated control samples after 7 days of culture (statistically proved by H(7)=22.451, P=0.002). Moreover, the depth of penetration rose substantially with increasing duration of culture for all cases, namely within the ranges of 50-70 μm, 80-90 μm and 90-120 μm (120 μm mostly for treated mats) from day 1 to day 7, respectively.

**Figure 8.**
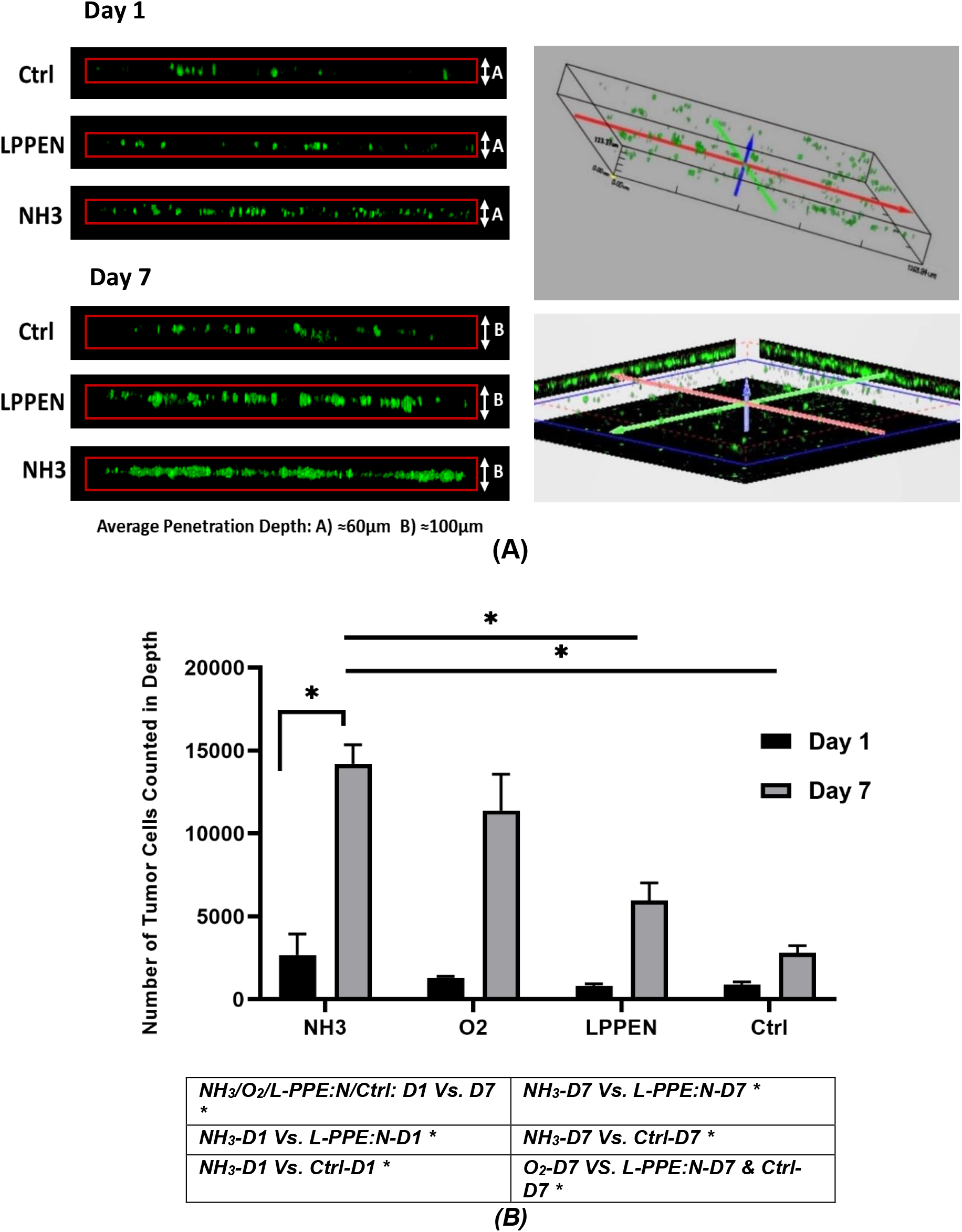
Tumor cell migration and growth in depth of the PP-3D-S model. (A): Representative 3D projections of stacked GFP breast cancer cells having migrated and proliferated through the depth of mats at day 1 and 7; (B) Number of GFP tumor cells quantified in depth of the 3D mat for different types of plasma treatments at day 1 and 7 (error bars: SE., n=3)

### 3.6. Drug screening experiments with PP-3D-S and Matrigel® models

Figure 9 shows images captured after 7 days, corresponding to increasing doses of Doxorubicin (Dox: 0, 0.05, 0.1, 0.5, 1 and 2 μM) with our PP-3D-S tumor interface model (see section 3.4.1). Treatment with increasing amounts of Dox are seen to have blocked GFP-tumor cell migration to the target tissue, fibroblasts in the 3D mat. We have compared this with invasion and migration of the same tumor cell types, along with the same Dox treatment regime, but now across Matrigel/Boyden chambers (images not shown). Quantitative data (histograms in Figure 9) show remarkably similar dose-dependent Dox-induced reductions of GFP-tumor cell migration in both models. Even though the two assays are clearly seen to have worked similarly, the PP-3D-S interface model is more advantageous because it can be set up far more rapidly (in roughly half the time), is adaptable to different tissue porosity and mechanics, and it avoids the characteristic batch-to-batch variation for which Matrigel^®^ is known.

**Figure 9.**
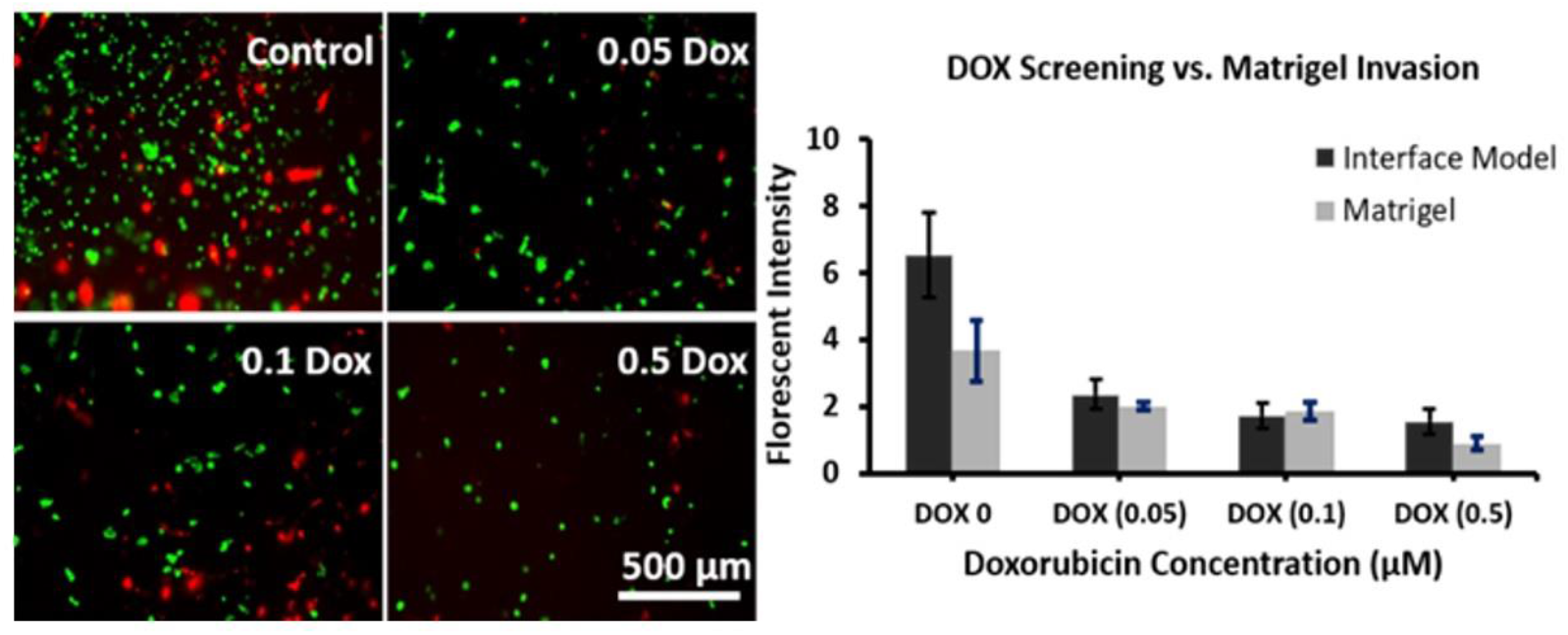
Images captured 7 days after treatment with increasing doses of Doxorubicin (Dox, μM) in our PP-3D-S tumor interface model. **Graph:** Comparison with invasion and migration of the same tumor cells across Matrigel/Boyden chambers (images not shown), showing similar Dox-induced reduction of GFP-tumor cell migration in both models (error bars: SD., n=3)

## 4. General Discussion and Conclusions

In this article we have reported results obtained with novel plasma-treated electro-spun 3D scaffolds combined with hydrogel (“*PP-3D-S*”) that can mimic various tissue types for general cell-biological research, including the human cancer microenvironment. This has significant potential value, including for customized cancer therapeutic screening; as a typical example of *PP-3D-S* use, designed to evaluate tumor cell migration, a breast-cancer tissue model has been simulated in this particular study. In more general terms, *PP-3D-S* has been found to enable mimicking “natural” cell interactions in a synthetic 3D environment that resembles the morphology of natural tissues in the human body, both in surface texture and chemistry as they affect the response by either benign or malignant living cells. It is also found to improve cell seeding and -adhesion capability of multi-cellular evaluation in 3D; controllable durability of the scaffold for predetermined duration, and improved controllable mechanical properties compared with commercially available 3D cell culture media such as Matrigel®. Mechanical characteristics can be made to vary over a wide range, to simulate softer or harder tissue types through choice of the electro-spun polymer scaffold, as demonstrated here by PLA, PCL and PU. PP-3D-S enables selecting the appropriate plasma treatment technique, either surface modification by grafting new functional groups (e.g. oxygen- or nitrogen-containing ones), or by coating the polymer fibre surfaces with thin plasma polymer films, both of which improve cell adhesion and -proliferation.

Regarding those plasma-induced surface treatments, the penetration of short-lived highly reactive species to large depths in the 3D scaffold, possibly exceeding 1000 μm, occurs thanks to their large mean-free-path lengths, especially in the case of the low-pressure plasmas used here. The beneficial result, of course, is uniform surface-chemical composition, hence uniform cell response throughout the scaffold volume. As clearly demonstrated, cell adhesion on plasma-modified mats greatly exceeds that on untreated controls. An unexpected and so far unexplained result has been the fact that L-PPE:N coatings only marginally improved performance with respect to the controls. Let us recall, first, that thin plasma polymer deposits *a priori* have an advantage over plasma modification (e.g. via O_2_, NH_3_, …), namely ability to avoid “hydrophobic recovery” (also known as aging), and second that L-PPE:N has performed very well in our past experience with a rather similar situation[44, 46, 47]; there is therefore a strong incentive to understand and correct this problem. This is already in progress in our laboratories at this time, via new atmospheric-pressure plasma-based coating approaches and chemistries, and preliminary results appear very promising indeed.

The purpose of developing the PP-3D-S co-culture system was to reproduce or mimic the physiological metastatic tumor microenvironment. The generated migration/invasion model could then be used to incorporate patient derived tumor cells and matching stromal fibroblasts for screening personalized therapies. The driving force for this migration across the interface between the top (hydrogel) and bottom (electrospun) scaffold is directly related to the surface treatment (Figure 5), but may also be related to signalling among the two different cell types [64-66]. In fact, combining fibroblasts as stromal cells co-cultured with tumor cells in the 3D interface model leads to an appropriate biochemical environment and more accurately represents realistic tissue environment around the tumor cells; this is essential for drug therapy experiments described in the presented work. Ongoing studies in our group are seeking to address specific aspects relating to the *ab-initio* presence of fibroblasts and ensuing signaling pathways among the different cell types within the co-culture 3D model, including patient-derived cells.

In two different sets of cancer drug screening experiments, a) *PP-3D-S* and b) Matrigel^®^, numbers of migrated surviving tumor cells were compared after exposure to varying dosages of the well-known chemotherapeutic Doxorubicin; numerical outcomes, ca. 75% reduction with 0.5μM drug concentration, were found to be very similar. These data indicate that *PP-3D-S* is an effective, economical and easy-to-use alternate 3D tumor migration model compared to standard Matrigel^®^ assays. Furthermore, it supports the notion that the PP-3D-S system represents a physiological tumor microenvironment model. This has implications for studying different cancer types, both primary and metastatic, as well as being tailored to a high-throughput format. In such a system, both academic and industrial labs may find valuable use with this model. Future work will test the PP-3D-S system using cancer cells of other origins (ie, prostate, lung and colo-rectal), or breast cancer cells, like here, but of differing degrees of aggressiveness, co-culturing patient-derived tumor cells and matching stomal cells, and a battery of alternate chemotherapeutics.

With the aim to provide a cell culture system of still lower cost, greater durability (shelf life) and reproducibility compared with existing commercial ones, we have carried out further research, concentrating on medium-sized fiber PLA scaffolds, but attempting to optimize other fabrication variables, including the mentioned atmospheric-pressure plasma treatments. Future work will probe all of these aspects.

## Acknowledgments

The authors gratefully acknowledge funding support from the Natural Sciences and Engineering Research Council (NSERC) of Canada (A.A. and M.R. W.), the Institute TransMedTech at Polytechnique Montreal (A.A., M.R.W. and D.H.R.), and Start-Up Funds from the Research Institute of McGill University Health Center and Fonds de Recherche du Quebec Sante (FRQS) Junior 2 Research Scholar Award (D.H.R.). The authors also gratefully acknowledge Professor Morag Park (McGill university) for kindly providing the fluorescent cell lines used in this study.

## Section of Supplementary Data

**Table** Process parameters applied to fabricate electrospun mats

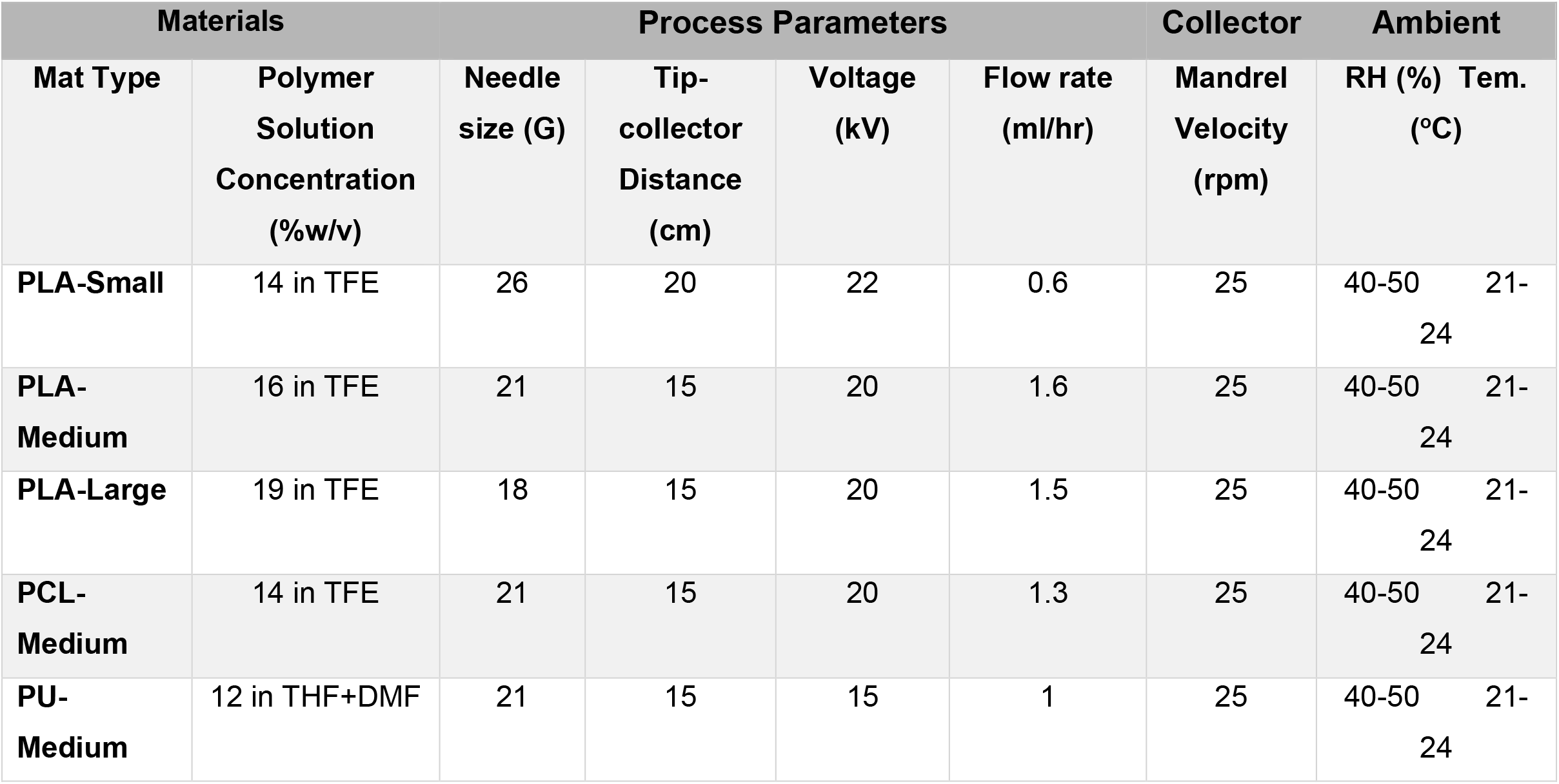

